# *Curtobacterium glycinis* sp. nov. from *Glycine max, Curtobacterium gossypii* sp. nov. from *Gossypium hirsutum* and *Curtobacterium oryzae* sp. nov. from *Oryza sativa*, three new *Curtobacterium* species and endophytes from agricultural crops

**DOI:** 10.1101/2021.03.18.434777

**Authors:** Sarah Seaton, Jacqueline Lemaire, Patrik Inderbitzin, Victoria Knight-Connoni, James F. White, Martha E. Trujillo

## Abstract

Three new *Curtobacterium* species from healthy tissues of agricultural crop plants in the United States are reported. They are *Curtobacterium glycinis* sp. nov. from soybean in Missouri, *Curtobacterium gossypii* from cotton in Puerto Rico and corn in Missouri, and *Curtobacterium oryzae* sp. nov. from rice in Texas.

## INTRODUCTION

Bacterial endophytes of plants have many beneficial interactions with their hosts and play a crucial role in promoting plant health (Finkel et al. 2017; Schenk et al. 2012). Microbiomes of the major agricultural crops including Cotton (*Gossypium hirsutum* L.), rice (*Oryza sativa* L.) and soybean (*Glycine max* (L.) Merr.) have been characterized (Edwards et al. 2015; Liu et al. 2019; Longley et al. 2020; Qiao et al. 2017; Roman-Reyna et al. 2020; Ullah et al. 2019), and beneficial endophytes have been detected (Bertani et al. 2016; de Almeida Lopes et al. 2018; Zhou et al. 2018). Plant-associated microorganisms are increasingly used for biotechnological applications, including biological control of plant pathogens, plant growth promotion, or isolation of active compounds (Ryan et al. 2008; Glick 2012; Bouizgarne 2013; Dey et al. 2014).

*Curtobacterium* is a genus of *Actinobacteria* in the family *Microbacteriaceae* comprising eight validly published species (Parte 2018). Several of the species have been isolated from agricultural plants. These include *Curtobacterium albidum* shown to counteract salt stress in rice (Vimal et al. 2019), *Curtobacterium flaccumfaciens* a pathogen of bean, beat, tulip and poinsetta (Collins and Jones 1983), *Curtobacterium herbarum* from grasses (Behrendt et al. 2002), and *Curtobacterium plantarum* from corn and soybean seed (Dunleavy 1989).

In this study, three *Curtobacterium* strains were isolated from healthy plant tissues in the United States. Strain OG107 from soybean in Missouri, strain VK105 from cotton in Puerto Rico, and strain SS108 from rice in Texas. The strains were characterized using molecular and phenotypic tests to determine their taxonomic placement. Our results indicate that the strains represent three new species of *Curtobacterium*, and we propose the names *Curtobacterium glycinis* sp. nov. (OG107), *Curtobacterium gossypii* sp. nov. (VK105) and *Curtobacterium oryzae* sp. nov. (SS108). These strains are taxonomically separated from the plant pathogens found within this genus.

## METHODS

### Isolation

Strains OG107 and SS108 were collected from the roots of healthy field-grown *Glycine max* in Missouri and *Oryza sativa* seedlings in Texas, United States, respectively. Plant tissue was washed with a mild detergent to remove particulates, surface-sterilized with bleach (1% v/v sodium hypochlorite) and ethanol (70% v/v), and homogenized. Serial dilutions of tissue homogenate were plated on a panel of media types for endophyte cultivation. Strain OG107, a small (0.7 mm diameter), pale yellow colony, arose on R2A agar after 5 days of incubation at 24°C, and strain SS108 arose as a non-descript colony on Actinomycete Isolation Agar. Both strains were streaked to purity and stored in glycerol (20% v/v) at −80°C until subjected to further testing.

Strain VK105 was isolated as described in Irizarry and White (2017). Briefly, seeds of wild, non-cultivated *G. hirsutum* plants were collected roadside in Guayama, Puerto Rico. Seeds were inoculated on Potato Dextrose Agar (PDA) and incubated at room temperature (25°C). Strain VK105 arose as an irregular yellow-orange pigmented colony. The colony was streaked to purity and stocked in 25% glycerol at −80°C until further analysis.

### Motility

Strains were tested for flagellar-dependent swimming and swarming motility on R2A plates solidified with 0.3% and 0.6% agar, respectively. Three independent colonies were inoculated onto R2A broth and grown for 36 hr at 24°C. Broth cultures were normalized to an OD600 of 0.1, and 1.5 μl of culture was spotted directly onto the surface of the motility agar. The diameter of colony expansion was measured for 5 days.

### Carbon source utilization

Substrate utilization was assessed using Biolog GenIII Microplates (Catalogue No. 1030) (Biolog Inc., Hayward, CA). Each bacterium was inoculated in duplicate plates using Protocol A described by the manufacturer, with the exception that plates were incubated at 30°C for 36 hr. Respiration leading to reduction of the tetrazolium indicator was measured by absorbance at 590 nm.

### Biochemical analyses

Catalase activity was evaluated by immediate effervescence after the application of 3 % (v/v) hydrogen peroxide solution via the tube method, a positive reaction was indicated by the production of bubbles. *Staphylococcus aureus* NCIMB 12702 and *Streptococcus pyogenes* ATCC 19615 were used as positive and negative controls, respectively. Oxidase activity was evaluated via the oxidation of Kovács oxidase reagent, 1% (w/v) tetra-methyl-p-phenylenediamine dihydrochloride in water, via the filter-paper spot method. A positive reaction was indicated when the microorganism’s color changed to dark purple. *Pseudomonas aeruginosa* NCIMB 12469 and *Escherichia coli* ATCC 25922 were used as positive and negative controls, respectively.

### Phylogenetic and genomic analyses

DNA was extracted from pure cultures using the Omega Mag-Bind Universal Pathogen Kit according to manufacturer’s protocol with a final elution volume of 60μl (Omega Biotek Inc., Norcross, GA). DNA samples were quantified using Qubit fluorometer (ThermoFisher Scientific, Waltham, MA) and normalized to 100 ng. DNA was prepped using Nextera DNA Flex Library Prep kit according to manufacturer’s instructions (Illumina Inc., San Diego, CA). DNA libraries were quantified via qPCR using KAPA Library Quantification kit (Roche Sequencing and Life Science, Wilmington, MA) and combined in equimolar concentrations into one 24-sample pool. Libraries were sequenced on a MiSeq using pair-end reads (2×200bp). Reads were trimmed of adapters and low-quality bases using Cutadapt (version 1.9.1) and assembled into contigs using MEGAHIT (version 1.1.2) (Li et al. 2015). Reads were mapped to contigs using Bowtie2 (version 2.3.4) (Langmead and Salzberg 2012), and contigs were assembled into scaffolds using BESST (2.2.8) (Sahlin et al. 2014).

Average nucleotide identity analyses were performed using the pyani ANIm algorithm (Richter and Rosselló-Móra 2009) implemented in the MUMmer package (Kurtz et al. 2004) retrieved from https://github.com/widdowquinn/pyani.

16S rRNA gene sequences were extracted from genome assemblies using barrnap (Seemann 2019) and submitted to GenBank. Phylogenetic analyses based on the 16S rRNA gene were performed using FastTree (Price et al. 2010) with a General Time Reversible substitution model. Taxon sampling for each species is described in the respective phylogenetic tree figure legend.

Average nucleotide identity (ANI) analyses between genome assemblies were performed using the pyani ANIm algorithm (Richter and Rosselló-Móra 2009) implemented in the MUMmer package (Kurtz et al. 2004) retrieved from https://github.com/widdowquinn/pyani.

Geographic distribution and host range of novel species were inferred by ANI to assemblies from unidentified species from GenBank (Ciufo et al. 2018) and the Indigo internal collection. An ANI threshold of ≥95% indicated conspecificity (Chun et al. 2018; Richter and Rosselló-Móra 2009). Distribution maps were generated in R version 4.0.4 (R Core Team 2021) with tigris 1.5 (Walker 2021), sf 1.0-2 (Pebesma 2018), ggplot2 3.3.2 (Wickham 2016), cowplot 1.1.1 (Wilke 2020) and other packages.

## RESULTS

### Phylogenetic and genomic analyses

#### *Curtobacterium glycinis* sp. nov. strain OG107

Strain OG107 shared 98.9% 16S rRNA gene sequence identity with *Curtobacterium citreum* DSM 20528^T^ and less with the remaining *Curtobacterium* species. A phylogenetic tree using FastTree (Price et al. 2010) confirmed the affiliation of strain OG107 with the genus *Curtobacterium*. OG107 was most closely related to *C. herbarum* P 420/07^T^ with 84% bootstrap support (Figure 1). The top average nucleotide identity (ANI) value of OG107 was 89.2% with *C. luteum* ATCC 15830^T^. This value was well below the threshold for species demarcation (Richter and Rosselló-Móra 2009; Chun et al. 2018) providing further genomic support that strain OG107 represents a new genomic species of *Curtobacterium*.

**Figure 1.**
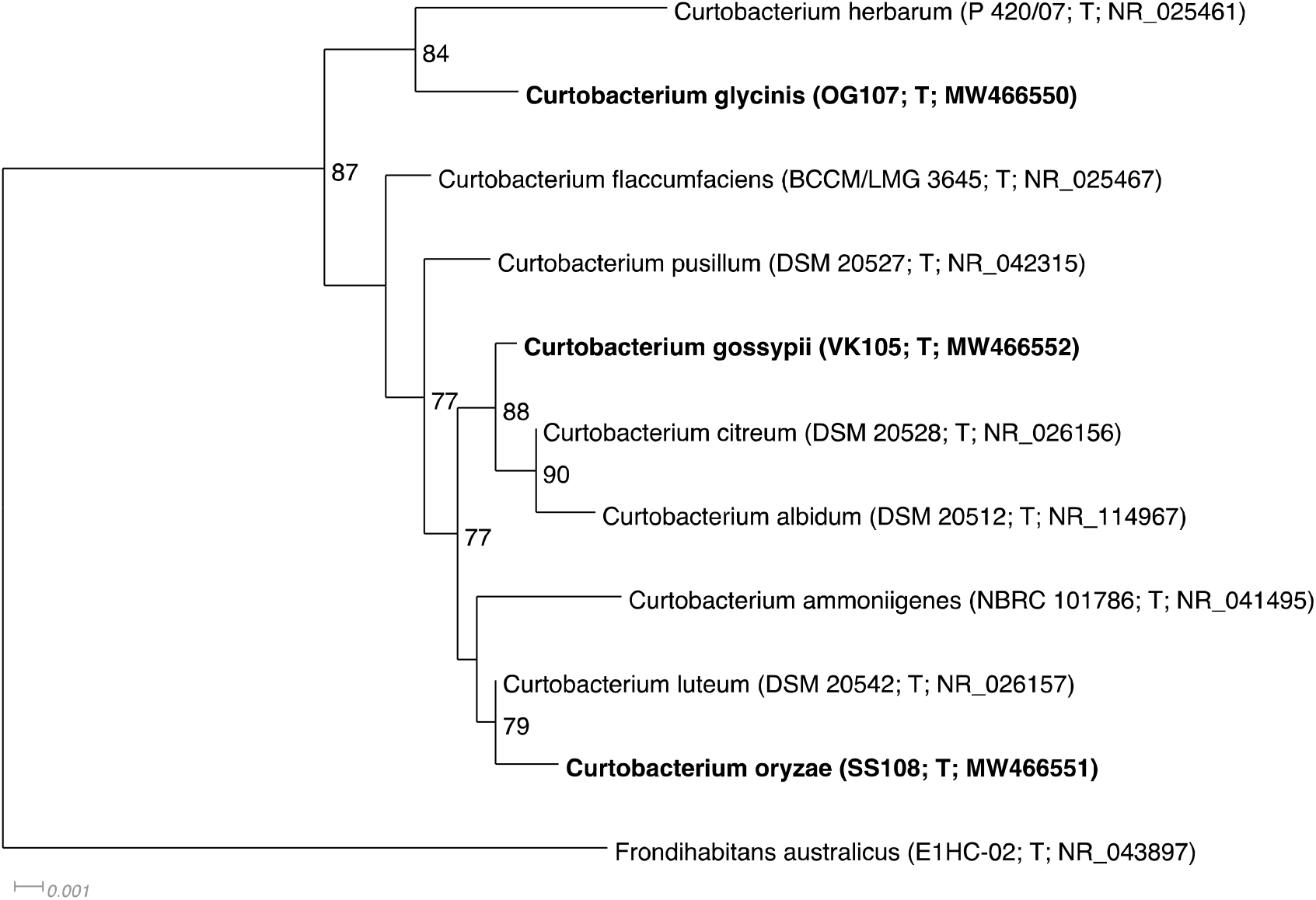
Phylogenetic 16S rRNA gene tree of *Curtobacterium glycinis* sp. nov. strain OG107, *Curtobacterium gossypii* sp. nov. strain VK105, *Curtobacterium oryzae* sp. nov. strain SS108 and relatives generated using FastTree (Price et al. 2010). All validly published *Curtobacterium* species were included in the tree, the tree is rooted with *Frondihabitans australicus*. New species are in bold. Strain identifiers and GenBank accession numbers follow species names, T stands for ‘type’. Support values above 70% are given by the branches. *Curtobacterium glycinis* is most closely related to *C. herbarum* with 84% support, *Curtobacterium gossypii* to *C. albidum* and *C. citreum* with 82% support and *Curtobacterium oryzae* to *C. luteum* with 79% support. Branch lengths are proportional to the changes along the branches, a scale bar is provided.

#### *Curtobacterium gossypii* sp. nov. strain VK105

Strain VK105 shared 99.8% 16S rRNA gene sequence identity with *Curtobacterium citreum* DSM 20528^T^ and less with the remaining *Curtobacterium* species. A phylogenetic tree using FastTree (Price et al. 2010) confirmed the affiliation of strain VK105 with the genus *Curtobacterium*. VK105 formed a monophyletic group with the species *C. albidum* and *C. citreum* supported by high bootstrap support (Figure 1) and was equally related to either of the two species. Average nucleotide identity (ANI) values of *C. albidum* DSM 20512^T^ and *C. citreum* DSM 20528^T^ to VK105 were both 87.7%. These values are well below the threshold for species demarcation (Richter and Rosselló-Móra 2009; Chun et al. 2018) providing further genomic support that strain VK105 represents a new genomic species of *Curtobacterium*.

#### *Curtobacterium oryzae* sp. nov. strain SS108

Strain SS108 shared 99.6% 16S rRNA gene sequence identity with *Curtobacterium luteum* strain DSM 20542^T^ and less with the remaining *Curtobacterium* species. A phylogenetic tree using FastTree (Price et al. 2010) confirmed the affiliation of strain SS108 with the genus *Curtobacterium*. SS108 was most closely related to *C. luteum* with 79% bootstrap support (Figure 1). The top average nucleotide identity (ANI) value of SS108 was 89.2% with *C. luteum* DSM 20542^T^. This value was well below the threshold for species demarcation (Richter and Rosselló-Móra 2009; Chun et al. 2018) providing further genomic support that strain SS108 represents a new genomic species of *Curtobacterium*.

### Geographic distribution and host range

Geographic distribution and host range of the novel species was inferred by comparison to congeneric genome assemblies from the Indigo internal collection. Hits included the Indigo strain *Curtobacterium gossypii* strain JL114 (ANI: 99.99%; query coverage: 99.33%) collected from healthy *Zea mays* plants in Missouri. Known geographic distributions of the novel species from culturing is illustrated in Figure 2, Figure 3 and Figure 4, and substrates are compiled in Table 1.

**Figure 2.**
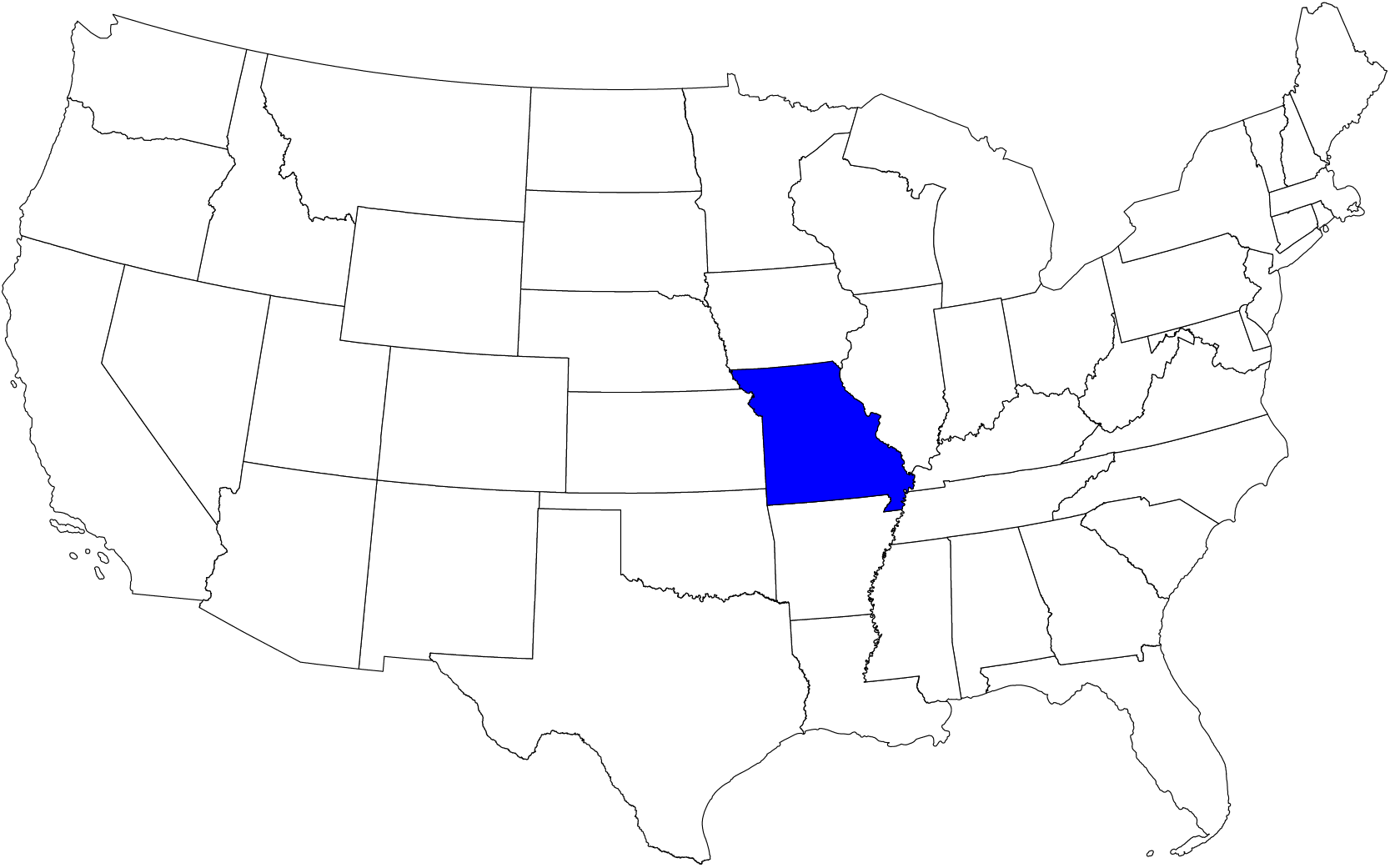
Geographic distribution of *Curtobacterium glycinis* sp. nov. based on culturing. Dark blue indicates state of origin for type strain.

**Figure 3.**
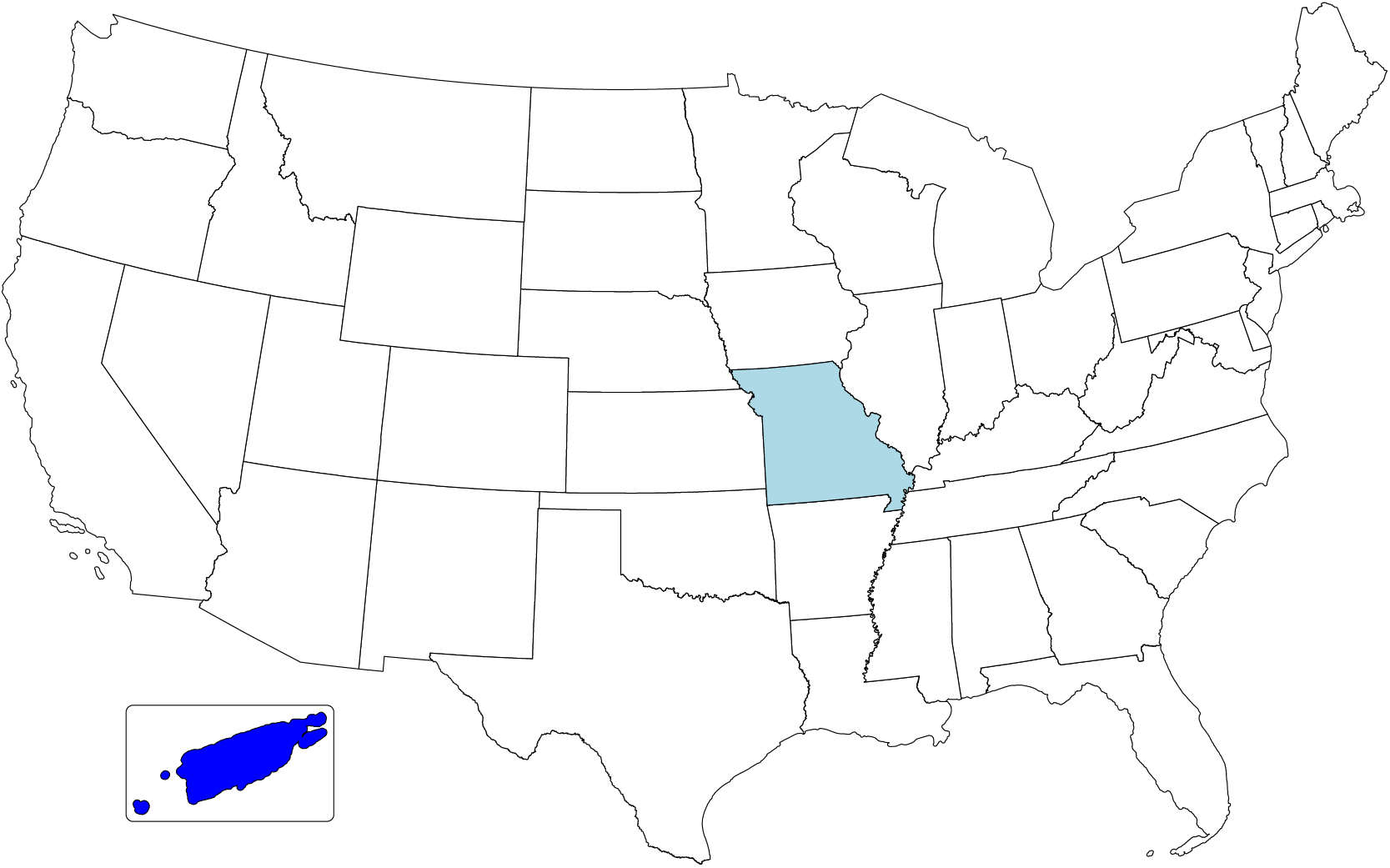
Geographic distribution of *Curtobacterium gossypii* sp. nov. based on culturing. Dark blue indicates state of origin for type strain, light blue additional strain. Inset: Puerto Rico (not to scale).

**Figure 4.**
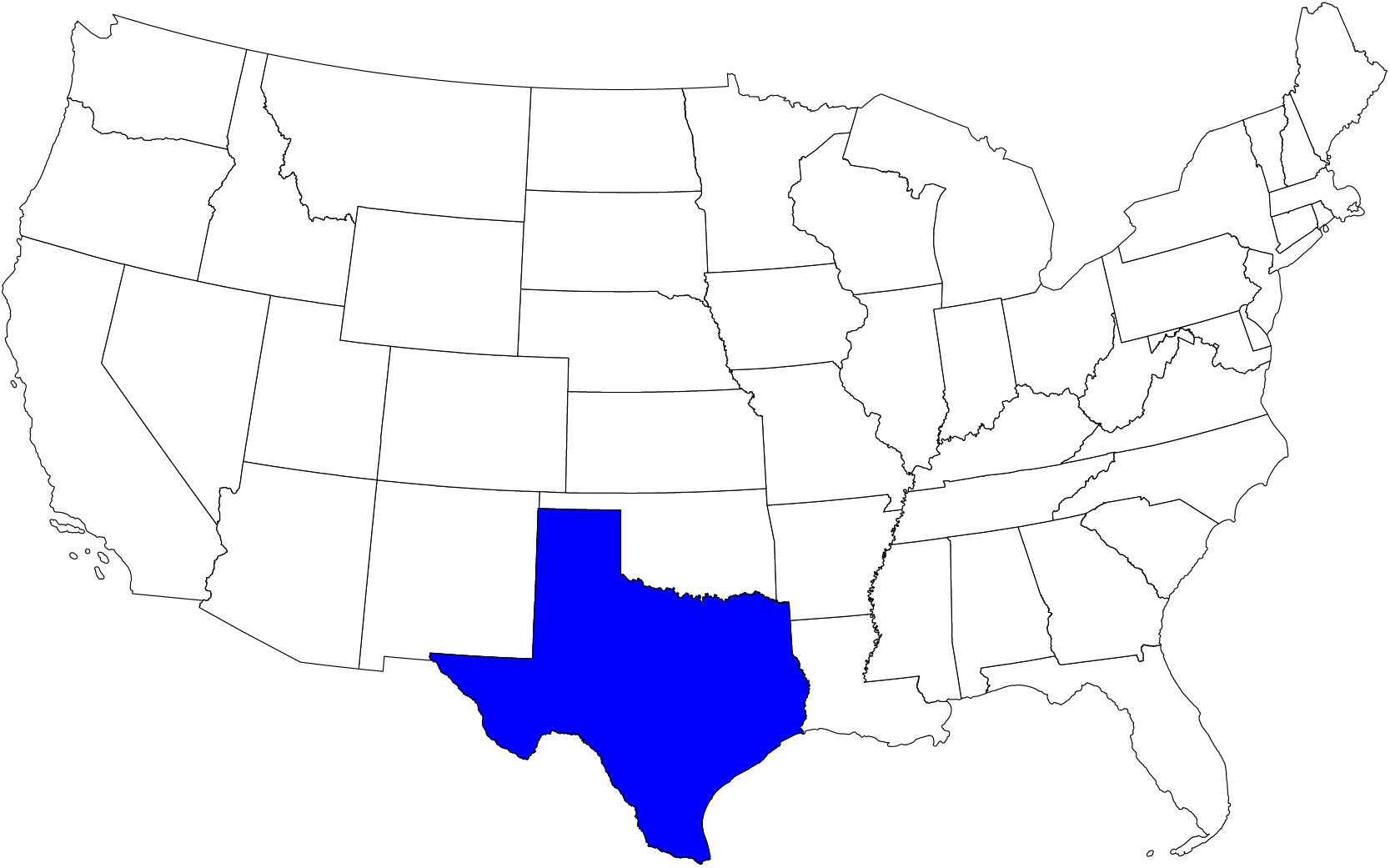
Geographic distribution of *Curtobacterium oryzae* sp. nov. based on culturing. Dark blue indicates state of origin for type strain.

**Table 1.** Substrates of novel species based on culturing.

| Species | Substrate<sup>1</sup> |
| --- | --- |
| *Curtobacterium glycinis* | *Glycine max* |
| *Curtobacterium gossypii* | *Gossypium hirsutum*, *Zea mays* |
| *Curtobacterium oryzae* | *Oryza sativa* |

<sup>1</sup> No disease symptoms reported for strains isolated from plants.

### Morphology, physiology and biochemical characteristics

#### *Curtobacterium glycinis* sp. nov. strain OG107

Strain OG107 stained Gram-positive. Cells had a slightly curved rod shape (0.2-0.4 μm in width and 1-3 μm in length); V and Y shaped forms were observed (Figure 5). They were aerobic, non-sporulating and showed motility. On nutrient agar, colonies were cream, entire, punctiform and convex. After 48 hr, good growth is observed on TSA, R2A and NA at 22 and 30°C. Growth was poor to moderate on the same media and incubation times at 37°C. Growth is observed when media are supplemented with 4% NaCl (w/v). Strain OG 107 was positive for catalase, but not for oxidase or urease. Being aerobic, the strain has an oxidative metabolism and according to the Biolog system, it is capable of using a range of substrates as carbon sources that include sugars, alcohol sugars and organic acids.

**Figure 5.**
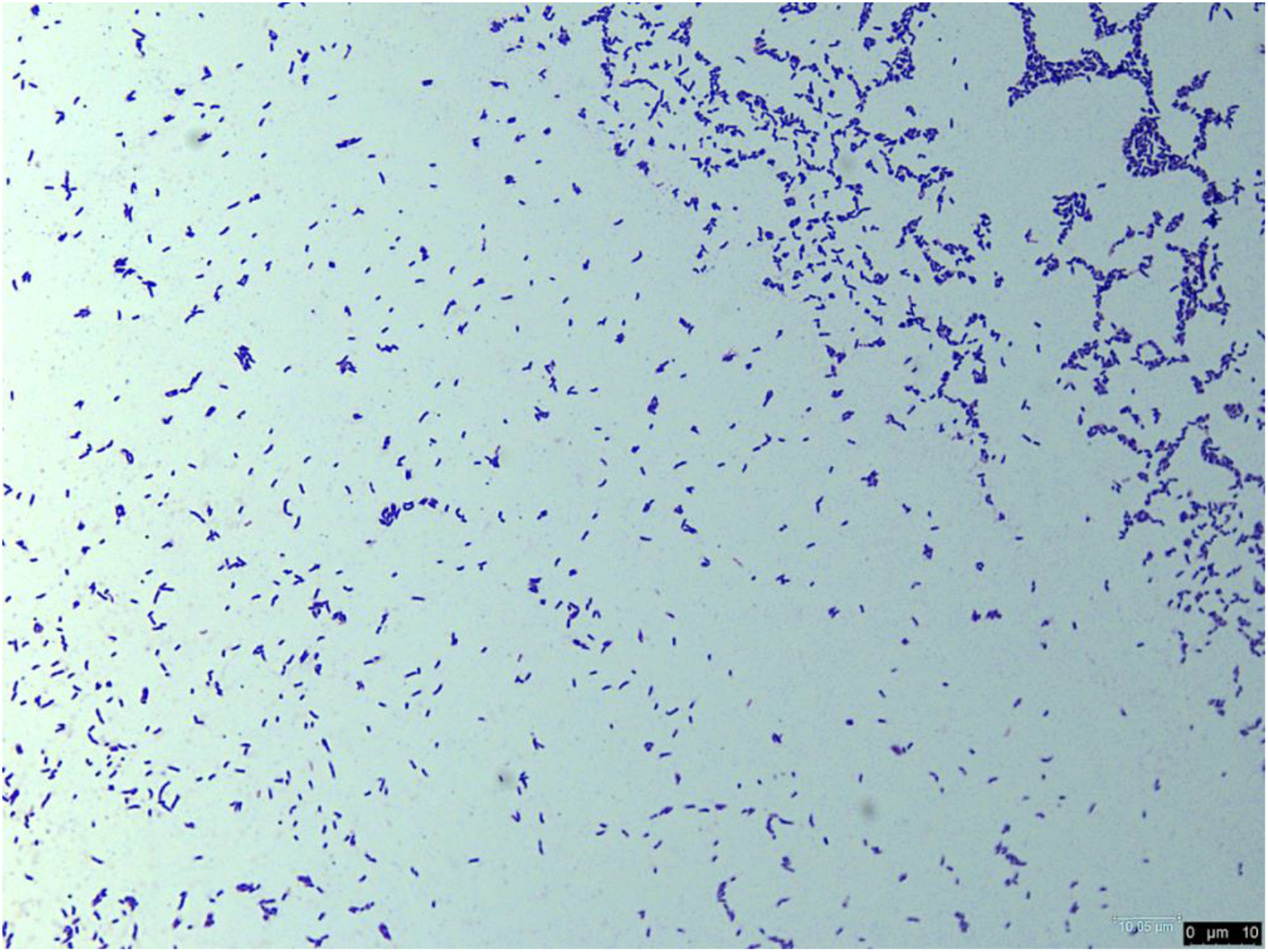
Morphology of *Curtobacterium glycinis* sp. nov. strain OG107 depicted following Gram stain using bright field microscopy. Bar = 10 μm.

Compared to its closest phylogenetic neighbor, *C. citreum* JCM 1345^T^, OG17 uses N-Acetyl-D-Galactosamine, D-arabitol, gentobiose, *myo*-inositol, mannitol and turanose as carbon sources. These tests help to differentiate between the two strains. A detailed description is given in Table 2 and the species description.

**Table 2.** Physiological characteristics of newly described *Curtobacterium glycinis, C. gossypii* and *C. oryzae* species and related *Curtobacterium* type strains. All strains were positive for the assimilation of dextrin, D-fructose, D-galactose, α-D-glucose, D-mannose and sucrose (data not shown).

|  | *C.*<br>*gossypii*<br>VK105 | *C.*<br>*oryzae*<br>OG107 | *C.*<br>*glycinis*<br>SS108 | *C.*<br>*citreum*<br>JCM<br>1345<sup>T</sup> | *C.*<br>*pusillum*<br>JCM<br>1350<sup>T</sup> | *C.*<br>*luteum*<br>JCM<br>1480<sup>T</sup> | *C.*<br>*albidum*<br>NBRC<br>15078<sup>T</sup> | *C.*<br>*herbarum*<br>DSM<br>14013<sup>T</sup> |
| --- | --- | --- | --- | --- | --- | --- | --- | --- |
| **General characteristics:** |  |  |  |  |  |  |  |  |
| **Colony pigment** |  | light yellow | ivory | yellow | yellow | yellow | ivory | orange |
| **Motility** | - | w | w | + | + | + | - | + |
| **Oxidase** | - | - | - | - | - | - | - | - |
| **Catalase** | + | + | + | + | + | + | + | + |
| **Urease** | - | - | - | - | - | - | - | nr |
| **Growth with 4% NaCl** | + | + | + | nr | nr | nr | nr | nr |
| **Growth with 8% NaCl** | + | - | - | nr | nr | nr | nr | nr |
| **Utilization of:** |  |  |  |  |  |  |  |  |
| **N-Acetyl-D-Glucosamine** | + | + | + | + | - | + | w | nr |
| **N-Acetyl-D-Galactosamine** | - | + | + | - | - | - | - | nr |
| **N-Acetyl Neuraminic Acid** | - | + | - | nr | nr | nr | nr | nr |
| **D-Arabitol** | + | + | + | - | w | - | - | nr |
| **D-Cellobiose** | + | + | + | + | - | + | + | nr |
| **L-Fucose** | w | + | + | + | - | + | - | nr |
| **Gentiobiose** | + | + | + | - | + | - | + | nr |
| **myo-Inositol** | + | + | + | - | - | - | - | nr |
| **α-D-Lactose** | + | + | + | + | - | + | + | nr |
| **D-Mannitol** | + | + | + | - | + | - | w | + |
| **D-Melibiose** | + | + | + | + | + | + | w | + |
| **β-Methyl-D-Glucoside** | + | + | + | - | + | - | w | nr |
| **D-Raffinose** | + | + | + | - | - | - | + | nr |
| **D-Sorbitol** | w | + | - | - | + | - | + | + |
| **Turanose** | + | + | + | - | + | - | + | nr |
| **D-Gluconic Acid** | + | + | + | - | + | - | + | nr |
| **D-Glucuronic Acid** | + | + | + | - | - | - | - | nr |
| **L-Alanine** | + | + | + | - | - | - | w | nr |
| **L-Glutamic Acid** | + | + | w | + | - | - | - | nr |
| **L-Rhamnose** | w | + | + | - | - | - | - | + |
| **D-Glucose-6-PO4** | - | + | + | - | - | - | - | nr |
| **Acetic Acid** | w | + | - | - | - | - | - | nr |

+, Positive; w, Weakly positive; -, Negative; nr, Not reported.
Data from this study, Aizawa et al. (2007) and Behrendt et al. (2002).

#### *Curtobacterium gossypii* sp. nov. strain VK105

The cells of strain VK105 stained Gram-positive, formed short irregular rods (0.4-0.6 μm in width and 2-4 μm in length), with cells diving by bending (Figure 6). Y shaped-forms were sometimes visible. Cells were non-motile and did not produce spores. Strain VK105 showed good aerobic growth on R2A and TSA agars. On both media, yellow to orange colonies were observed after 2 days incubation at 22 and 30°C. At 7 days, colonies were circular, smooth, glistening and slightly convex. The new strain produced catalase but not oxidase. Strain VK105 has an oxidative metabolism and uses a range of substrates as carbon sources that are also useful for to differentiate against its closest phylogenetic neighbors, *Curtocabterium citreum* and *C. albidum*, namely D-arabitol, L-fucose, D-glucuronic acid, and *myo*-inositol. These and other physiological characteristics are presented in Table 2 and the species description.

**Figure 6.**
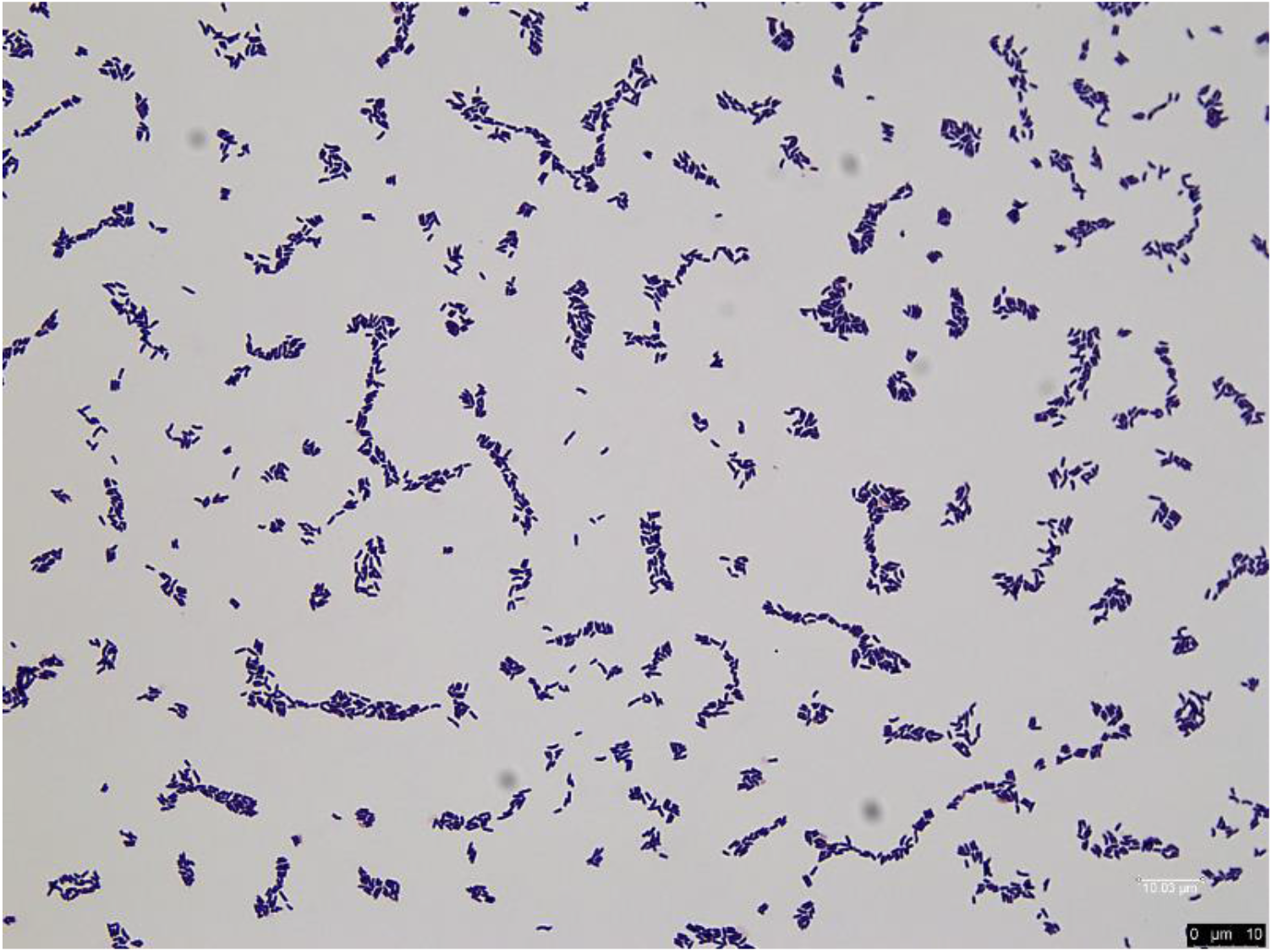
Morphology of *Curtobacterium gossypii* sp. nov. strain VK105 depicted following Gram stain using bright field microscopy. Bar = 10 μm.

#### *Curtobacterium oryzae* sp. nov. strain SS108

Cells stained-Gram positive, are non-motile, non-spore-forming and rod shaped or slightly curved. In many cases, two cells remained together and formed V shapes (Figure 7). Average cell size ranged from 0.2-0.4 μm in width and 1.5 × 2 μm in length. Colonies of strain SS108 were light yellow, entire, round and raised when grown on nutrient agar. After 2 days, good growth was observed on TSA and R2A agars at 22, 30 and 37°C. On nutrient agar, good growth was obtained at 30°C after 2 days and was moderate at 22 and 37°C. However, after 7 days, good growth was also seen at 22°C. Aerobic growth.

**Figure 7.**
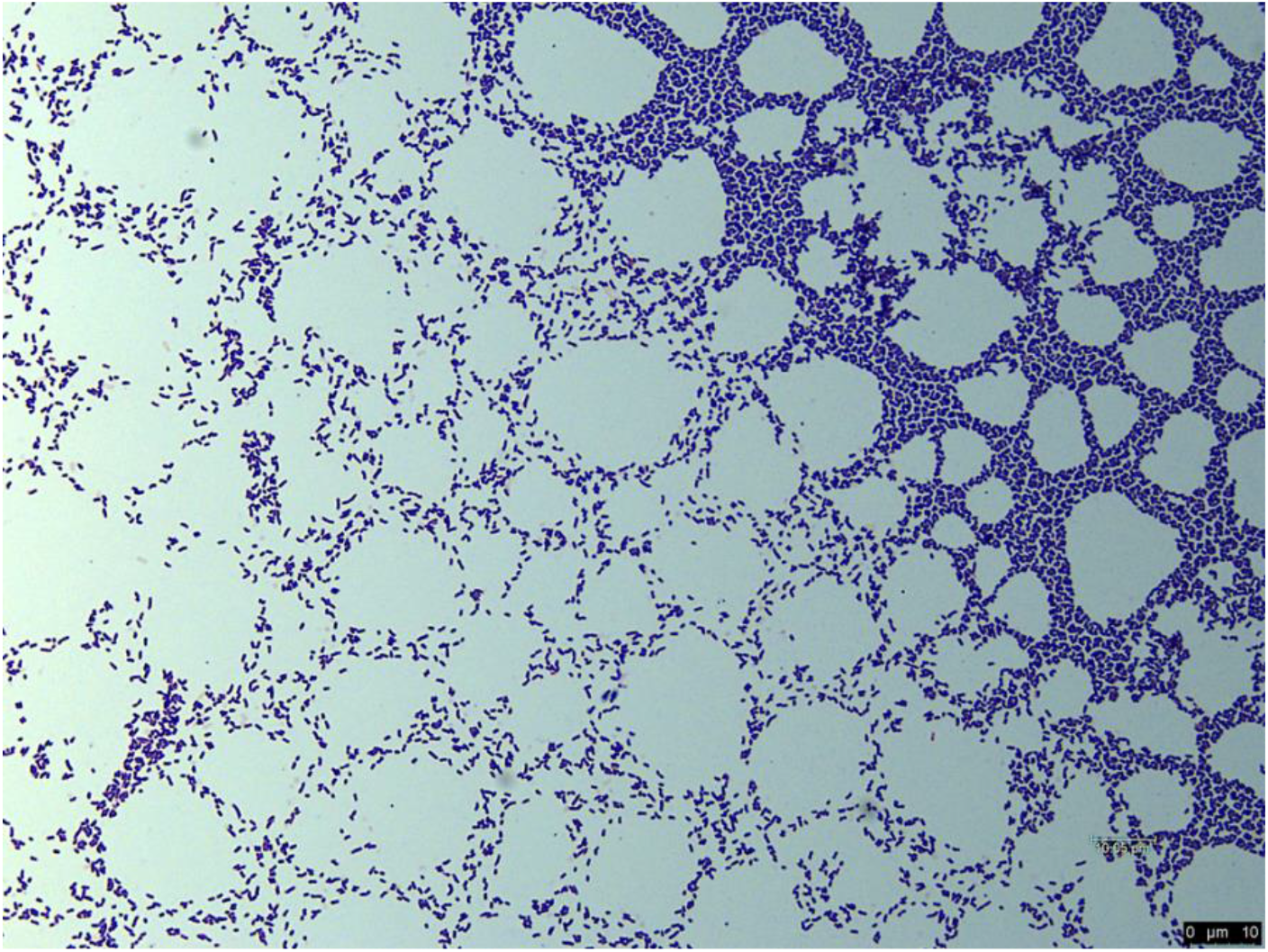
Morphology of *Curtobacterium oryzae* sp. nov. strain SS108 depicted following Gram stain using bright field microscopy. Bar = 10 μm.

Tolerates 4% NaCl (w/v). Enzyme activity was detected for catalase, but not for oxidase or urease. Several carbon sources are useful to differentiate between the new strain and *C. luteum* JCM 1480^T^ and include the utilization of N-Acetyl-D-Galactosamine, D-arabitol, gentobiose, *myo*-inositol and D-mannitol among others (Table 2). Other phenotypic characteristics are given in the species description.

## DESCRIPTION OF CURTOBACTERIUM GLYCINIS SP. NOV

*Curtobacterium glycinis* (gly.ci’nis. N.L. fem. gen. n. *glycinis* of *Glycine max*, the soybean, referring to the origin of the type strain)

Cells are Gram-stain positive, non-spore forming, motile and rod-shaped with V and Y shapes produced. Aerobic and chemoorganotrophic. Colonies are circular, smooth, glistening and slightly convex. On R2A and nutrient agar, colonies are cream color and entire, punctiform and convex. Growth is abundant at 22 and 30°C but moderate at 37°C. Grows in the presence of 4% NaCl. Growth is observed between pH 5-7. Positive for catalase; negative for oxidase and urease. The following substrates are used as carbon sources: N-acetyl-D-glucosamine, N-acetyl-D-galactosamine, D-arabitol, D-cellobiose, L-fucose, gentiobiose, *myo*-inositol, α-D-lactose, D-mannitol, D-melibiose, β-methyl-D-glucoside, D-raffinose, turanose, D-gluconic acid, D-glucuronic acid, L-alanine, L-glutamic acid (weak), L-rhamnose, D-Glucose-6-PO4.

The type strain is resistant to nalidixic acid and aztreonam.

The type strain OG 17^T^ was isolated from the roots of healthy field-grown *Glycine max* in Missouri, USA.

## DESCRIPTION OF CURTOBACTERIUM GOSSYPII SP. NOV

*Curtobacterium gossypii* (gos.sy’pi.i. N.L. gen. n. *gossypii* of *Gossypium*, the generic name of cotton, referring to the origin of the type strain)

Cells are Gram-stain positive, non-motile, non-spore-forming, coccoid or rod-shaped. Aerobic and chemoorganotrophic. Yellow to orange colonies on R2A and TSA agars. Growth on these media is observed after 48 h. Catalase is produced but not oxidase. Grows in the presence of 1 - 4 % NaCl; weak at 8% and at pH 5-7. The following substrates are used as sole carbon sources: dextrin, D-maltose, D-trehalose, D-cellobiose, gentiobiose, sucrose, D-turanose, stachyose, D-raffinose, α-D-lactose, D-melibiose, β-methylD-glucoside, D-salicin, N-acetyl-D-glucosamine, α-D-glucose, D-mannose, D-fructose, D-galactose, inosine, sodium lactate, D-mannitol, D-arabitol, *myo*-inositol, glycerol, L-aspartic acid, L-glutamic acid, pectin, D-arabitol, D-gluconic acid, D-glucuronic acid and glucuronamide.

The type strain is resistant to nalidixic acid and aztreonam.

The type strain, VK105^T^, was isolated from seeds of wild cotton collected in Puerto Rico, USA.

## DESCRIPTION OF CURTOBACTERIUM ORYZAE SP. NOV

*Curtobacterium oryzae* (o.ry′zae. N.L. gen. n. *oryzae* of rice, referring to the origin of the type strain)

Cells are Gram-stain positive, non-motile, non-spore-forming and rod shaped or slightly curved; V shapes are formed. Aerobic and chemoorganotrophic. Colonies are light yellow, entire, round and raised on nutrient agar. Good growth is obtained at 22, 30 and 37°C on the media tested. Grows on media supplemented with 4% NaCl and at pH 5-7.

Positive for catalase, but negative for oxidase or urease. The following substrates are used as carbon sources: N-Acetyl-D-Glucosamine, N-Acetyl-D-Galactosamine, N-Acetyl Neuraminic Acid, D-Arabitol, D-Cellobiose, L-Fucose, Gentiobiose, myo-Inositol, α-D-Lactose, D-Mannitol, D-Melibiose, β-Methyl-D-Glucoside, D-Raffinose, D-Sorbitol, Turanose, D-Gluconic Acid, D-Glucuronic Acid, L-Alanine, L-Glutamic Acid, L-Rhamnose, D-Glucose-6-PO4 and Acetic Acid.

The type strain is resistant to nalidixic acid and aztreonam.

The type strain, SS108^T^, was isolated from *Oryza sativa* seedlings collected in Texas, USA.

## ACKNOWLEDGEMENTS

We would like to thank Professor Aharon Oren, The Hebrew University of Jerusalem, for checking Latin species names. Support was provided by USDA-NIFA Multistate Project W4147 and the New Jersey Agricultural Experiment Station.

